# Photon-separation to enhance the spatial resolution in pulsed STED microscopy

**DOI:** 10.1101/408286

**Authors:** Giorgio Tortarolo, Yuansheng Sun, Kai-Wen Teng, Yuji Ishitsuka, Luca Lanzanó, Paul R. Selvin, Beniamino Barbieri, Alberto Diaspro, Giuseppe Vicidomini

## Abstract

Stimulated emission depletion microscopy (STED) is one of the pivotal super-resolution techniques. It overcomes the spatial resolution limit imposed by the diffraction by using an additional laser beam, the STED beam, whose intensity is directly related to the achievable resolution. Despite achieving nanometer resolution, much effort in recent years has been devoted to reduce the STED beam intensity because it may lead to photo-damaging effects. Exploring the temporal dynamics of the detected fluorescence photons and accessing the encoded spatial information has proven to be a powerful strategy, and has contributed to the separation by lifetime tuning (SPLIT) technique. The SPLIT technique uses the phasor analysis to efficiently distinguish photons emitted from the center and the periphery of the excitation spot. It thus improves the resolution without increasing the STED beam intensity. This method was proposed for architectures based on STED beam running in continuous wave (CW-STED microscopy). Here, we extend it to architectures based on pulsed STED beam (pSTED microscopy). We show, through simulated and experimental data, that the SPLIT-pSTED method reduces the detection volume of the pSTED microscope without significantly reducing the signal-to-noise ratio of the final image, thus effectively improving the resolution without increasing the STED beam intensity.

## 1 Introduction

Fluorescence microscopy has established as an invaluable tool for the study of life sciences, thanks to features such as non-invasiveness, molecular specificity and sensitivity. Furthermore, the introduction of super-resolution microscopy techniques has allowed to overcome the spatial resolution limit imposed by diffraction,^1^ and thus to investigate biological phenomena with unprecedented resolution.^2^ Stimulated emission depletion (STED) microscopy was one of the firstly introduced super-resolution techniques;^3,4^ the typical point scanning STED microscope co-aligns the conventional Gaussian excitation beam with the so called STED beam, engineered to generate a donut-shaped intensity distribution at the focus, i.e., with a ”zero”-intensity point in the center. The STED beam forces all fluorophores, except those in the ”zero”-intensity point at the center of the excitation spot, to de-excite to the ground state *via* stimulated emission, thus reducing the region from which the fluorescence signal is emitted and then recorded. Albeit theoretically the resolution of STED microscopy can reach the molecular size by increasing the intensity of the STED beam, practically it is limited by other factors, such as the noise^5^ and the amount of laser power that can be delivered to the sample in order to prevent photo-damage effects.^6^ For the latter reason, much effort has been spent on reducing the peak power of the STED beam necessary to reach a given spatial resolution. One turnkey insight has been the comprehension of the variations in fluorescence temporal dynamics induced by the STED beam itself. The stimulated emission process opens a new fluorophore’s de-excitation pathway, whose rate (instantaneous probability) strongly depends on the intensity of the STED beam. Since the STED beam intensity is spatially distributed as a donut, the position of the fluorophore with respect to the center of the excitation spot is encoded in it’s fluorescence temporal decay (slower at the center of the excitation spot and faster at the donut crest). This understanding has been exploited to reduce the peak intensity of the STED beam needed to achieve a certain resolution, enabling effective sub-diffraction resolution also in such conditions where the STED beam intensity is not sufficient to obtain a complete fluorescent depletion/quenching.^7–9^

The first application of this principle is the so-called gated-CW-STED microscope.^8,10^ If the STED beam is implemented with a continuous-wave (CW) laser, the peak intensity reduces and the effective resolution as well. However, the farther is the fluorophore from the focal point, the shorter is its (perturbed) excited state lifetime *τ*_*ST ED*_ (i.e., the average time that the fluorophore spend in to the excited state); thus, by using a pulsed excitation and a time-gated detection scheme - fluorescence is registered after a delay from the excitation events - the fluorescence at the periphery due to the incomplete depletion is removed and the resolution improves. On the other hand, time-gating rejects also a portion of ”wanted” photons from the center of the focal spot, resulting in a reduced signal-to-noise/background ratio of the final image that may cancel out the resolution enhancement.^5,9^ For implementations based on pulsed STED beam, the benefits of exploring the fluorescent temporal dynamics and of the time-gated detection depend on the pulse-width of the STED beam.^9^ Early pulsed STED (pSTED) microscopes used pulse-width below 200 ps, thus much shorter than the excited-state of typical organic fluorophores (*τ*_*fl*_ *∼* 1-10 ns). This temporal condition makes the fluorescence emitted during the action of the STED beam and the incomplete depletion negligible, thus the time-gated detection useless.^11^ More recently it has been shown that STED microscopy based on sub-nanosecond (*∼* 600-1000 ps) pulsed lasers substantially reduces the photo-bleaching compared to early pSTED implementations:^12^ photo-bleaching is supra-linear with the STED beam intensity.^13,14^ In this case, since the pulse-width is comparable with the excited-state lifetime *τ*_*fl*_ and the peak intensity reduces (for a given average intensity of the pulse), the fluorescence emitted during the action of the STED beam is not anymore negligible and the benefit of time-gating is relevant. For these reasons, most of the current pSTED microscopy implementations - including the commercial systems - relies on sub-nanosecond fiber laser and implement a time-gated detection (gated-pSTED microscopy). However, similarly to gated-CW-STED microscopy, also for gated-pSTED microscopy the SNR of the final image is reduced.

*A-posteriori* approaches, such as multi-image deconvolution^15^ and separation of photons by life-time tuning (SPLIT),^16^ can solve this problem. In particular, the SPLIT method analyses the pixel fluorescent temporal decays within the phasor framework, and represents a straightforward approach to separates all the photons emitted from the long lifetime fluorophores located in the focal point (”wanted” photons), from the ”unwanted” photons emitted from the short lifetime fluorophores located in the focal periphery. Here, we extend the SPLIT method to pulsed STED microscopy (SPLIT-pSTED), allowing to recover the resolution hidden by the incomplete depletion/quenching, but without impacting on the SNR.

A first method based on the phasor analysis of a pSTED images has been developed by us^17^ and successively improved by Wang L. et al.^18^ This method uses the phasor-plot representation – of the pSTED image – to implement a pixel segmentation on the raw pSTED image that selects the pixels characterized by a slow fluorescence temporal dynamics, thus composed primarily by ”wanted” photons, and discards the pixels characterized by a fast fluorescence temporal dynamics, thus composed primarily by ”unwanted” photons. On the contrary, the SPLIT approach selects from each pixel only the ”wanted” photons, rather than performing a simple binary pixel classification. In essence, the SPLIT-pSTED method that we propose – compared to the phasor-plot based segmentation methods – sorts photons and not pixels, thus providing more quantitative and artifact-free pSTED images. In the context of quantitative imaging, it is important to highlight that the SPLIT imaging technique preserves the linearity in the image, i.e., the pixel intensity values are linear with the fluorophore concentration.^16^

## 2 Materials and Methods

### Phasor-based analysis

The SPLIT method is based on the analysis of the fluorescent signal dynamics by means of the so-called phasor analysis. The phasor analysis is a powerful tool able to describe the evolution of a signal (as a function of a variable, such as the time) as a single point in a plane with coordinates *g* (or cosine transform) and *s* (the sine transform): the phasor plot. In the context of SPLIT-STED imaging, the phasor approach is performed on every pixel of a time-resolved measurement (e.g., on the histogram of the photon-arrival times associated to any pixel) to discriminate molecular species with different temporal fingerprints. In particular, the phasor analysis is used to discriminate the photons emitted by the fluorophores localized in the center of the STED detection volume from the photons emitted by the fluorophores localized in the peripheral region, because these two classed of fluorophores are characterized by different excited-state lifetime, thus fluorescent decay dynamics. In brief, it can be used to further shrink the detection volume, thus improving the effective spatial resolution of a STED microscope.

We start by considering a single fluorophore and a CW-STED microscopy architecture, i.e., the STED beam runs in CW. Since the Gaussian-shaped excitation and the donut-shaped depletion intensity foci are co-aligned, if the fluorophore is in the very center of the excitation spot (*r* = 0), it does not interact with STED (or stimulating) photons, and thus it emits fluorescence according to its unperturbed lifetime *τ*_*fl*_. The decay dynamics of such fluorophore (thus, the histogram of the photon-arrival times) may be described with a single exponential exp(*-t/τ*_*fl*_) and corresponds to a single point in the phasor space, that lies on the semicircle of radius 1*/*2 with center (1*/*2, 0) (Fig. 1). On the contrary, if the fluorophore is located in the ring described by 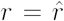, it interacts with STED photons and is likely to be quenched to the ground state. However, as a result of the non perfect efficiency of the stimulated emission process, it emits fluorescence nonetheless, but following a faster single exponential decay law 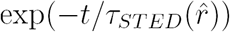, where 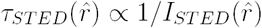 and *I*_*ST ED*_(*r*) is spatial (radial symmetric) intensity distribution of the STED beam. The corresponding point in the phasor space is still laying on the same semicircle, but shifted toward higher *g* values, the limiting case of infinite STED intensity is the point (1, 0) (Fig. 1).

**Fig 1:**
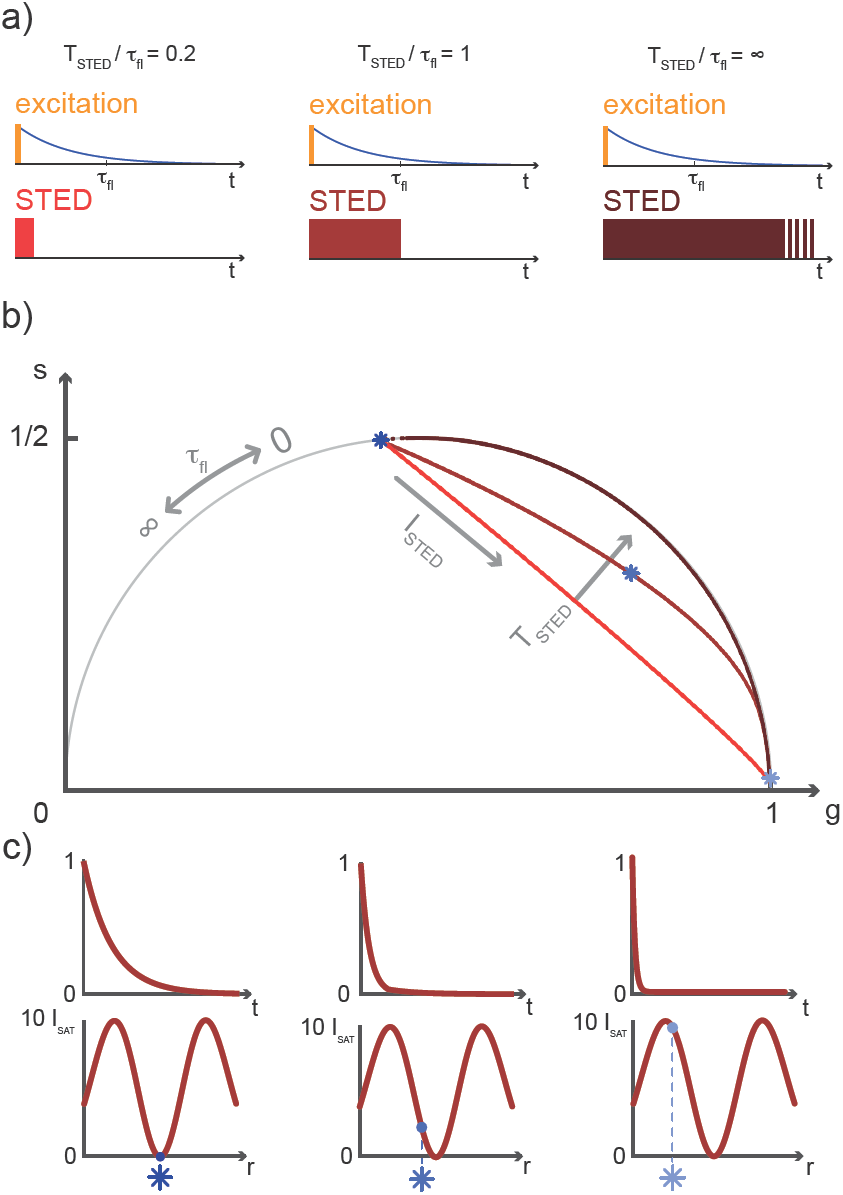
Phasor plot trajectories of a single molecule exposed to increasing STED beam intensities, for different STED beam pulse-widths. (a) Three cases of increasing *γ* = *T*_*ST ED*_*/τ*_*fl*_ are represented, where *T*_*ST ED*_ is the STED beam pulse-width and *τ*_*fl*_ the unperturbed fluorescence lifetime: *γ* = 0.2 (red), *γ* = 1 (deep red) and *γ* = *∞* (garnet), the continuous-wave STED laser configuration. (b) Synthetic pSTED experiment of a single molecule exposed to increasing doses of stimulating photons, for the three pulse-widths cases depicted in (a). All trajectories in the phasor space obtained increasing the STED intensity start from the point in the semicircle describing the unperturbed fluorescence lifetime *τ*_*fl*_, and end in the point (*g, s*) = (1, 0) describing instantaneous decay. If *γ* = 0.2 (very short STED pulse-width, red), the trajectory is a chord in the semicircle; in the limit case of *γ* = *∞* (CW configuration, deep red), the trajectory is an arc of the semicircle. Values of *γ ∈* (0.2, *∞*) leads to trajectories lying between the two limiting cases. (c) We now consider the condition *γ* = 1 (deep red trajectory): if the molecule lyes in the very center of the excitation spot (left, dark blue star) it does not interact with stimulating photons, thus its unperturbed temporal decay is represented in the phasors space as the corresponding point in the semicircle. Increasing the distance of the molecule from the center of the excitation spot leads to faster decays, due to higher STED intensities (center, blue star and right, light blue star): the corresponding points in the phasor space are thus closer to (*g, s*) = (1, 0). Repetition rate of the simulated excitation and STED beams: 60 MHz.

When considering a pulsed STED (pSTED) implementation, the key difference with the CW-STED implementation lies on the temporal dynamics of the fluorescence photons from the peripheral region of the excitation spot: the intensity decay dynamics of a fluorophore illuminated by the pulsed STED beam can be described with a piecewise function, characterized by a temporal decay that is faster during the STED pulse (0 *≤ t < T*_*ST ED*_), and slower when the STED beam is off (*t ≥ T*_*ST ED*_):

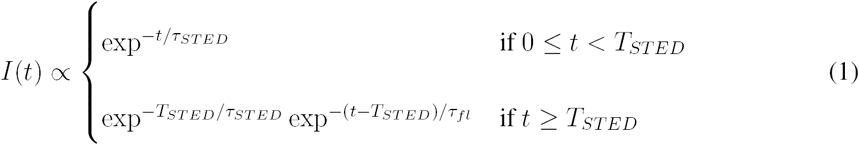

As a results, a fluorophore in the periphery of the excitation spot is represented in the phasor space by a point that is lying inside of the semicircle, conversely to the CW-STED implementation. In particular, this point is lying on a trajectory which again moves from the point associated to the unperturbed fluorophore (no stimulated photons) to the point (1, 0), for increasing STED intensity. Notably, the shape of the trajectory in a pSTED system dependents on the ratio between the pulse-width of the STED beam, *T*_*ST ED*_, and the natural lifetime of the fluorophore, *τ*_*fl*_, i.e., *γ* = *T*_*ST ED*_*/τ*_*fl*_. The two considered limit cases for *γ* are (i) very small pulse-width, *γ* = 0.2: the trajectory is a chord in the semicircle (Fig. 1b, red trajectory); infinite large pulse-width, *γ >>* 1 (equivalent to a CW-STED implementation): the trajectory is an arc of the semicircle (Fig. 1b, garnet red trajectory).

In a real STED imaging experiment the fluorescent signal registered by each pixel is the sum of the signals provided by all the fluorophores located within the detection region associated to the pixel. Thus, to reduce the effective detection region and improve the resolution without increasing the intensity beam, the phasor analysis isolates, from the fluorescent signal of each pixel, the component associated to the fluorophores located in the center of the excitation region, i.e., the slower components.

Although the dependency *τ*_*ST ED*_(*r*) yield to a continuous family of populations of emitters located at increasing distances from the center of the excitation spot and with increasingly faster temporal decays, for simplicity reasons it is convenient to divide them in two species: the first one related to the center of the excitation spot (species 1, **P**_1_ in the phasor space); the second one related to the periphery (species 2, **P**_2_ in the phasor space), that originates the undesired signal. Now, the problem becomes to isolate the component of the signal generated from the first species. As per the rules of phasors, a combination of such two species will be represented in the phasor space as a point lying on the line **P**_1_**P**_2_. As described in the SPLIT technique,^16^ the contributes from these two species can be separated *via* a linear decomposition approach in order to extract only the signal originated by fluorophores in the center of the excitation spot.

We now consider a time-resolved measurement. The total number of collected photons *N* in a given pixel is the sum of photons *N*_1_ and *N*_2_, emitted by molecules in the center and in the periphery of the excitation spot, respectively. As described before, the corresponding vector in the phasor space *P* can be described as the linear combination of the two vectors **P**_1_ = (*g*_1_, *s*_1_) and **P**_2_ = (*g*_2_, *s*_2_) describing the two pure species: **P** = (*N*_1_**P**_1_ + *N*_1_**P**_2_)*/N* = *f*_1_**P**_1_ + *f*_2_**P**_2_, where *f*_1_ and *f*_2_ are the fractional components of the detected photons. We can write this linear system in the matrix form **P** = M **f**, where **f** = (*f*_1_, *f*_2_) and *M* = (**P**_1_, **P**_2_) is the matrix containing the temporal dynamics of the two species in the phasor domain. The solution **f** = *M*^−1^**P** allows to separate photons emitted by molecules in the center of the excitation spot *N*_1_ = *f*_1_*N* from the undesired contributions of peripheral molecules *N*_2_ = *f*_2_*N*. The final image with higher resolution is obtained by iterating this process for each pixel of the raw time-resolved image.

It is important to observe that this phasor analysis is not restricted to simple single exponential decay, as for the CW-STED implementation, but it is valid in general for each signal evolution, thus also for the signal generated by the fluorophores in the case of p-STED microscopy.

### Pulsed-STED microscope

For this study we performed measurements using an ISS Alba confocal/STED laser scanning microscopy system (www.iss.com/microscopy/instruments/alba.html) coupled with a Nikon Te2000 microscope.^17^ The excitation source is a 640-nm picosecond pulsed diode laser (Becker Hickl, BDL-SMN-640, 120 ps pulse width). A 775-nm sub-nanosecond pulsed fiber laser (OneFive, Katana 775, 600 ps pulse width) provides the STED beam. The two lasers are synchronized by either (a) using the depletion laser at the 40 MHz repetition rate (master) to trigger the excitation laser (slave); or (b) using the excitation laser at the 50 MHz repetition rate (master) to trigger the depletion laser (slave). Both lasers are also synchronized to the FastFLIM module to perform time-resolved STED measurements. The 640-nm excitation laser is mounted on the ISS 3-diode laser launcher to control its intensity, and then delivered to the Alba system via a single mode polarization maintained fiber (QiOptics). The 775-nm STED laser intensity is controlled by the ISS intensity control unit consisting of a motorized rotating half-wave plate and a fixed Glan-Thompson polarizer. We use an optical delay line (custom made) for the fine tuning (picosends) of the temporal delay between the STED pulses with respect to the excitation beam; the STED laser is then delivered via a single mode PMF (Thorlabs) to the STED beam conditioning module (custom made), to generate the donut-shaped profile of the delpetion beam. Inside the Alba module, the excitation and the STED beams are combined using a 670 long-pass dichroic mirror (Semrock); we use a pair of galvanometric mirrors to scan both beams in the plane of the sample perpendicular to the optical axes. The objective is the highly numerical aperture Nikon Plan APO *λ* 60X/1.4NA oil. The scanning device is synchronized to the data acquisition unit (FastFLIM by ISS) for time-resolved Digital Frequency Domanin (DFD) measurements. We use another dichroic mirror (custom made by Chroma) to separate the descanned fluorescence signal from the excitation and STED light. We filter the emitted fluorescence by both a 720 nm short-pass NIR light blocking filter (Optical Density 8, Chroma), and a 679/41 nm band-pass emission filter (Semrock). We set the motorized confocal pinhole (from *∼*20 *μ*m to 1 mm) placed before the detector to 60 *μ*m in diameter (*∼*1 Airy unit) for STED measurements. The detection unit is the single photon counting module avalanche photodiode (SPCM-ARQH-15 by Excelitas). Data acquisition is performed using the ISS VistaVision 64-bit software; phasor analysis is performed by a custom written Matlab code, which is provided upon request.

### Sample preparation

*Fluorescent beads and nanorulers.* In this study we used crimson fluorescent beads of 60 nm diameter (FluoSpheres Red and Crimson), diluted in water 1:1000 (v/v) for a sparse sample. We dropped the diluted solution of fluorescent beads onto a poly-L-lysine (Sigma) coated glass coverslip and waited 5 min; then we washed the coverslip with water and dried it by blowing nitrogen onto it. Finally, we mounted it with a special medium (Mounting Medium, Invitrogen). *Fixed HeLa cells.* Human HeLa cells were transfected with plasmid DNA encoding GFP-ActA-Halotag^19^ (ActA binds to the outer mitochondrial membrane) overnight using Lipofectamine 2000 (Life Technologies) following manufacturers protocol. The cells were fixed using 4% paraformaldehyde (Fisher Scientific), and permeabilized with 0.2% Triton-X 100 (Sigma). The cells were then labeled with ATTO647-GBP (GFP-Booster, Chromotek) in the presence of 3% bovine serum albumin (BSA) in PBS. After labeling the cells were fixed using 4% paraformaldehyde for 15 minutes.

## 3 Results

### Synthetic data

We first applied the SPLIT method on the synthetic pSTED image of a single fluorophore. This simulation allows also to derive the point-spread-function (PSF) of the SPLIT-pSTED system (Fig. 2a). In this context, it is important to remember that the SPLIT-STED microscope is - in general - a linear and space-invariant system,^20^ thus its PSF fully describes the characteristic of the system, including the spatial resolution. Given the Gaussian-and donut-shaped focal intensity distributions of the excitation and depletion beams, respectively, and considering a STED pulse width of *T*_*ST ED*_ = 600 ps, we can calculate the expected fluorescence decay for each pixel of the image (Eq. 1), i.e. the so called temporal effective PSF of the pSTED microscope. In essence, the calculation of the temporal effective PSF is equivalent to simulate molecules located at different distances between the center of the excitation spot (*r* = 0), where the STED power is theoretically null, and *r*_*crest*_, where the STED intensity is maximized. Successively, for each pixel, we calculated the phasor values and we reported them in the phasor plot. The corresponding family of points in the phasor space describes a trajectory that: (i) since we simulated a realistic situation in which the center of the donut-is not a perfec ”zero”-intensity point, does not start from a point located on the semicircle but from a point (which represents the point *r* = 0) inside the semicircle; (ii) due to the limited intensity of the STED beam, the trajectory does not terminate in the limiting point (1, 0), but the latest point represents the position *r*_*crest*_, where the STED intensity is maximum (Fig. 2b). We then applied the linear decomposition described above. As *P*_1_ component (wanted signal, i.e., central signal), we chose the starting point of the trajectory, the one associated to the slowest dynamics, and as *P*_2_ (unwanted signal, i.e., peripheral signal) a point located at the tail of the trajectory. Results show that the method is effective in rejecting the contribution of fluorophores from the periphery of the excitation spot (Fig. 2, b), and thus in a reduction of the effective PSF (Fig. 2 b).

**Fig 2:**
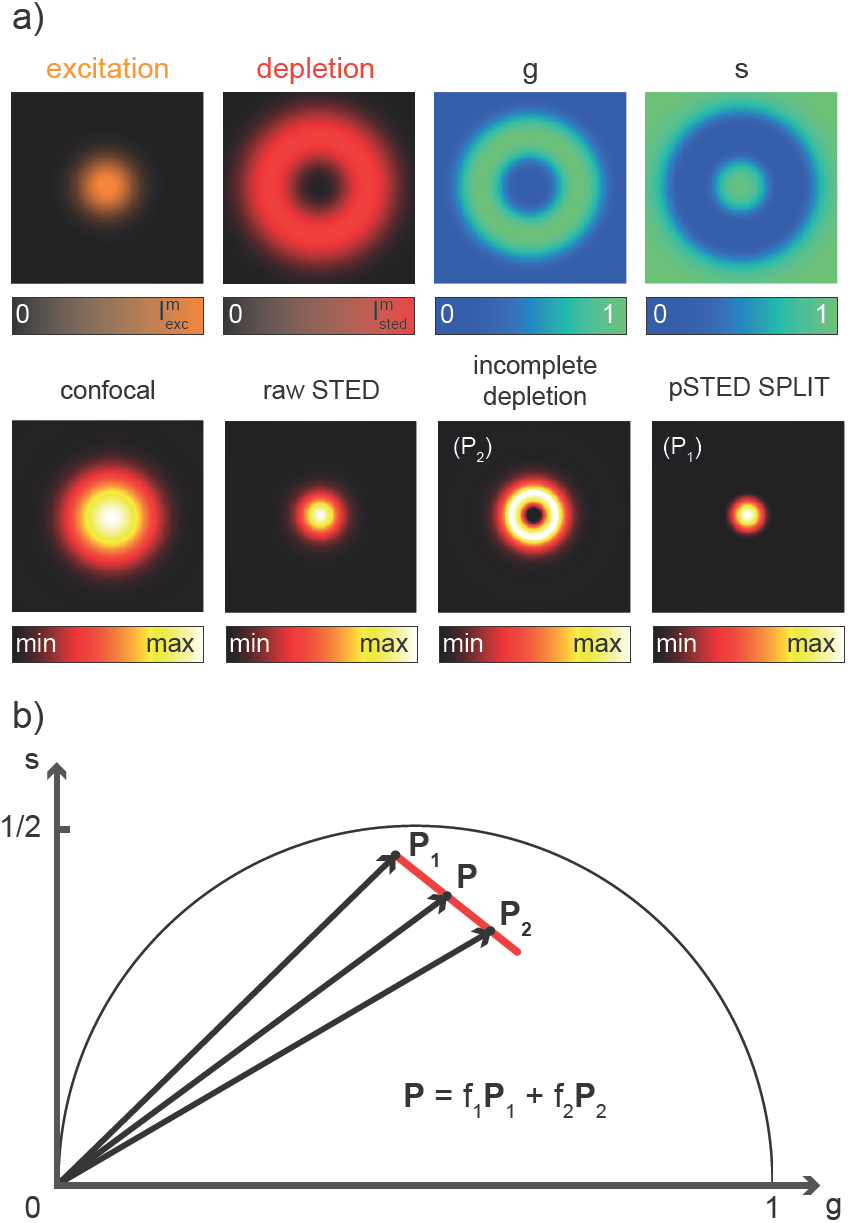
The pSTED-SPLIT method applied to synthetic data. (a) Simulation of the temporal effective point spread function (tE-PSF) of a pSTED experiment, equivalent to simulate the temporal dynamics of molecules located ad different distances *r* with respect to the center of the excitation spot. The *g* and *s* images of the rawSTED data indicate different temporal dynamics as a function of *r*. Notably, also for *r* = 0 (center of the excitation spot) the temporal decay of the molecule is perturbed, since the depletion intensity profile features a non-zero minimum to mimic real-life pSTED experiments. Thus, the point *P*_1_ representing such molecule in the phasor space (b) does not lye exactly on the semicircle. As expected, increasing *r* values are related to point closer to (*g, s*) = (1, 0). Given two points *P*_1_ and *P*_2_ related to central molecules and peripheral ones respectively, the pSTED-SPLIT method removes the fluorescence signal arising from the latter (a, incomplete depletion) *via* a linear decomposition approach. The result (a, pSTED SPLIT) shows a PSF that is shrunk when compared to the rawSTED counterpart.

### Real data

We successively validated the SPLIT-pSTED approach on real data obtained using phantom sample, such as fluorescent beads, and biological sample.

We first tested the performances of our STED setup by performing a series of time-resolved pSTED measurements with increasing depletion power, imaging a sample of 60 nm sized Crimson fluorescent beads (Figure 3). As expected, the cluster of points in the phasor plot related to all pixels in the image lyes close to the semicircle when the STED beam is off; as the depletion power increases and the spatial resolution of the resulting images improves, the cluster elongates toward the point (1, 0) following the simulated expected trajectory. The non perfect overlapping of the centroid of the cluster may be consequence of the high concentration of fluorophores on the beads, which induces a self-quenching phenomenum.^9^

**Fig 3:**
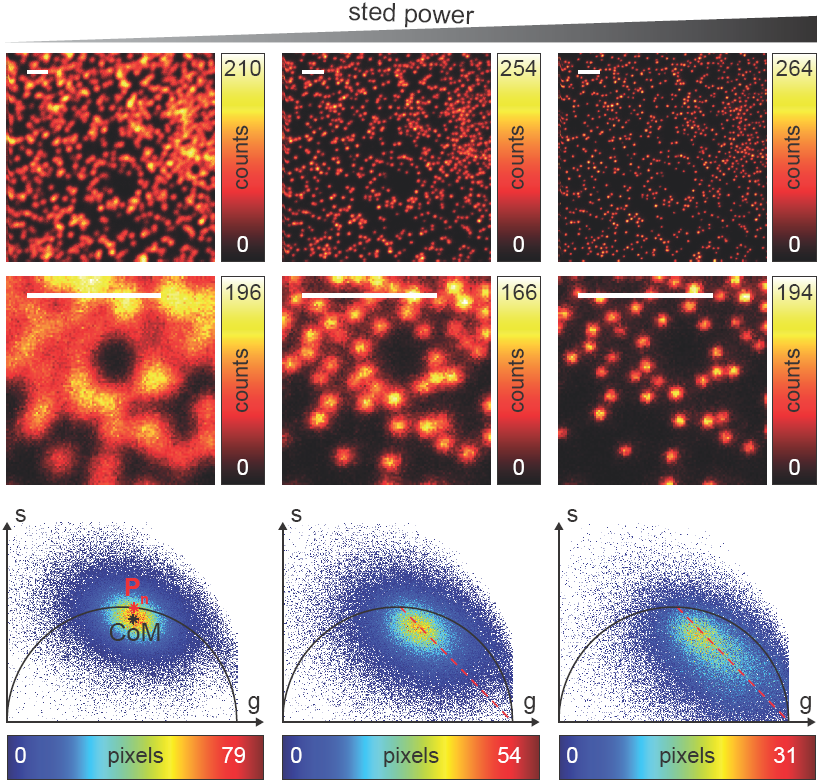
Time-resolved pSTED measurements of 60 nm sized fluorescent beads with increasing STED power. Images clearly reveal a resolution improvement when increasing the STED power, from confocal imaging (*P*_*ST ED*_ = 0 mW, left) to pSTED imaging (*P*_*ST ED*_ = 25 mW, center; and *P*_*ST ED*_ = 91 mW, right). Magnified views of the recorded images are shown in the central row. The phasor analysis (bottom) reveals the increasingly perturbed temporal dynamics of incomplete-depleted fluorophores, as expected from simulations (dotted red line). Pixel-dwell time: 100 *μ*s. Pixel-size: 20 nm. Format: 512 *×* 512 pixels. Scale bars: 1 *μ*m.

We then tested the linear decomposition approach on the same sample of 60nm sized crimson fluorescent beads. In real data a critical aspect of the SPLIT-pSTED is the choice of the two points **P**_1_ and **P**_2_ in the phasor plot. In this work we apply the following protocol for each measurement (Fig. 4 a,b): (i) perform a time-resolved confocal measurement (STED beam is off) to retrieve the unperturbed fluorescence lifetime of the fluorophore; fix **P**_*n*_ as the corresponding point on the semicircle at the shortest distance from the centroid (center of mass, CoM) of the cluster of points in the phasor plot; (ii) simulate via a custom-made Matlab tool a molecule/fluorophore with a lifetime described by **P**_*n*_; we then consider a STED impulse of duration *T*_*ST ED*_ = 600 ps, and calculate the different temporal behaviors of such molecule when interacting with increasing doses of stimulating photons: in the phasors space, the different decays are described by a family of points, or expected trajectory; (iii) perform a time-resolved pSTED measurement of the same area of step (i): the phasor analysis reveals a distribution of points in accordance with the expected trajectory; (iv) select both **P**_1_ and **P**_2_ on the simulated trajectory. A point **P**_1_ shifted towards the (1, 0) with respect to **P**_*n*_ allows to take into consideration the non-ideal ”zero” intensity condition; (v) force all the points **P**(*x, y*) to the line in between **P**_1_**P**_2_ and the expected trajectory; then find the fractional components **f**(*x, y*) = *M* ^-1^(*x, y*)**P**(*x, y*). We finally obtain the pSTED-SPLIT image as *N*_1_(*x, y*) = **f**_1_(*x, y*)*N* (*x, y*) (Fig. 4 c). Following this procedure the p-STED SPLIT approach succeeds in rejecting contributions from the periphery of the excitation spot and thus shrinks the size of the effective PSF. The reduced effective PSF leads to better resolved images when compared to the raw STED counterpart, i.e., the raw data collected during the time-resolved pSTED measurement, before applying the decomposition algorithm. Notably, the maximum counts of the raw and pSTED-SPLIT images are similar, which indicate that the SNR is not reduced during the process (Figure 4).

**Fig 4:**
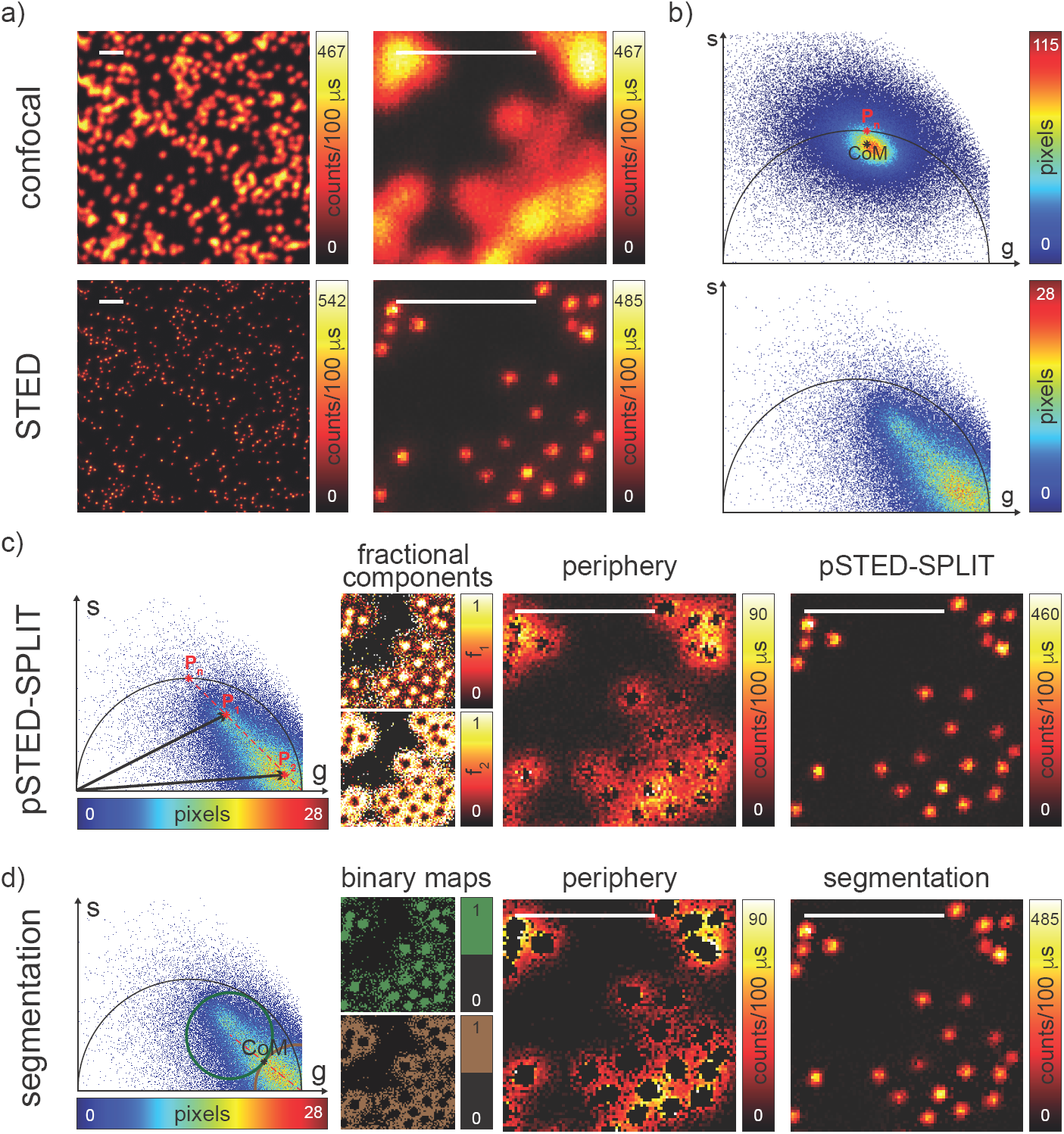
pSTED-SPLIT analysis of fluorescent beads. Results of the linear decomposition and of the pixel segmentation approaches for time-resolved pSTED measurements of 60 nm sized crimson fluorescent beads. (a) Resolution is clearly increasing from confocal imaging (top) to pSTED imaging (bottom); magnified views are also reported. The phasor analysis (b) shows how the STED beam perturbs the temporal dynamics of the fluorophores. (c) The pSTED-SPLIT method shown for magnified views of recorded data: points *P*1 and *P*_2_ are chosen from the phasor plot on the simulated trajectory (red dot line) calculated for point *P*_*n*_. For each pixel, ”wanted” and ”unwanted” photons are sorted thanks to the linear decomposition approach: the corresponding fractional components *f*1 and *f*2 allow to generate the final pSTED-SPLIT result and the image obtained solely by rejected photon, respectively. Notably, the signal to noise ratio of the raw pSTED image is preserved by the pSTED-SPLIT analysis. (d) The pixel segmentation approach: in the phasor space, we define a circular region of interest (RoI) centered in the limiting point *P*_*l*_ = (1, 0) passing through the center of mass (CoM, black star) of the distribution of points (brown circle). A second circular RoI with the same radius is defined to be tangent with the first one and with the center lying on the straight line identified by *CoM* -*P*_*l*_ (green circle). The two resulting binary maps are related to pixels dominated by peripheral and central fluorophores, respectively. The final segmentation result is then obtained by applying the binary map on the raw pSTED intensity image. Pixel-dwell time: 100 *μ*s. Pixel-size: 20 nm. Format: 512 *×* 512 pixels. Scale bars: 1 *μ*m.

At this stage it is also interesting to compare the SPLIT-pSTED approach with the phasor-based segmentation that we proposed in.^21^ In short, we use the CoM of the cluster of points in the phasor plot to generate two circular regions of interest with same radius and touching – externally tangent – in the cluster’s CoM. The first region – centered in the point (1, 0) – selects pixels with a shorter fluorescence temporal decay, thus composed primarily by ”un-wanted” photons. The second region selects pixels with a longer fluorescence temporal decay, thus composed primarily by ”wanted” photons. By back-projecting the pixels lying in the second region we obtained a segmented pSTED images (Fig. 4 d). Notably, the method proposed by^18^ better selects the two regions on the phasor plot (”abandoned” and ”selected” area), since it allows considering all the pixel contained in the semicircle, but similarly to our method the final pSTED result is a segmented version of the raw image. A close comparison of fluorescent beads imaging between the SPLIT method and the phasor-based segmentation method shows only a marginal improvement of the former with respect to the later. However, the results are substantially different for imaging of more convoluted structures, such as the mithocondria membrane, where the SPLIT-pSTED method shows superior performance. We compare the proposed SPLIT method with the phasor-based segmentation method on fixed HeLa cells with TOM20 labeled mithocondria (Figure 5). Thanks to the photons separation – rather than the pixel separation, the SPLIT-pSTED method is able to reveal the localization of the TOM20 proteins in the membrane of the mithocondria with higher spatial resolution (with respect to confocal) and without introducing artifacts. On the contrary, the phasor-based segmentation method tends to generate spotty structures, which may not fully represent the real morphology of the mithocondria membrane. Similarly to intensity-based segmentation methods – the phasor-based segmentation methods highlight the brighter pixels, which produce good-looking results when imaging well separated point-like structures, such as the fluorescent beads, but tends to lose morphological information for other structures.

**Fig 5:**
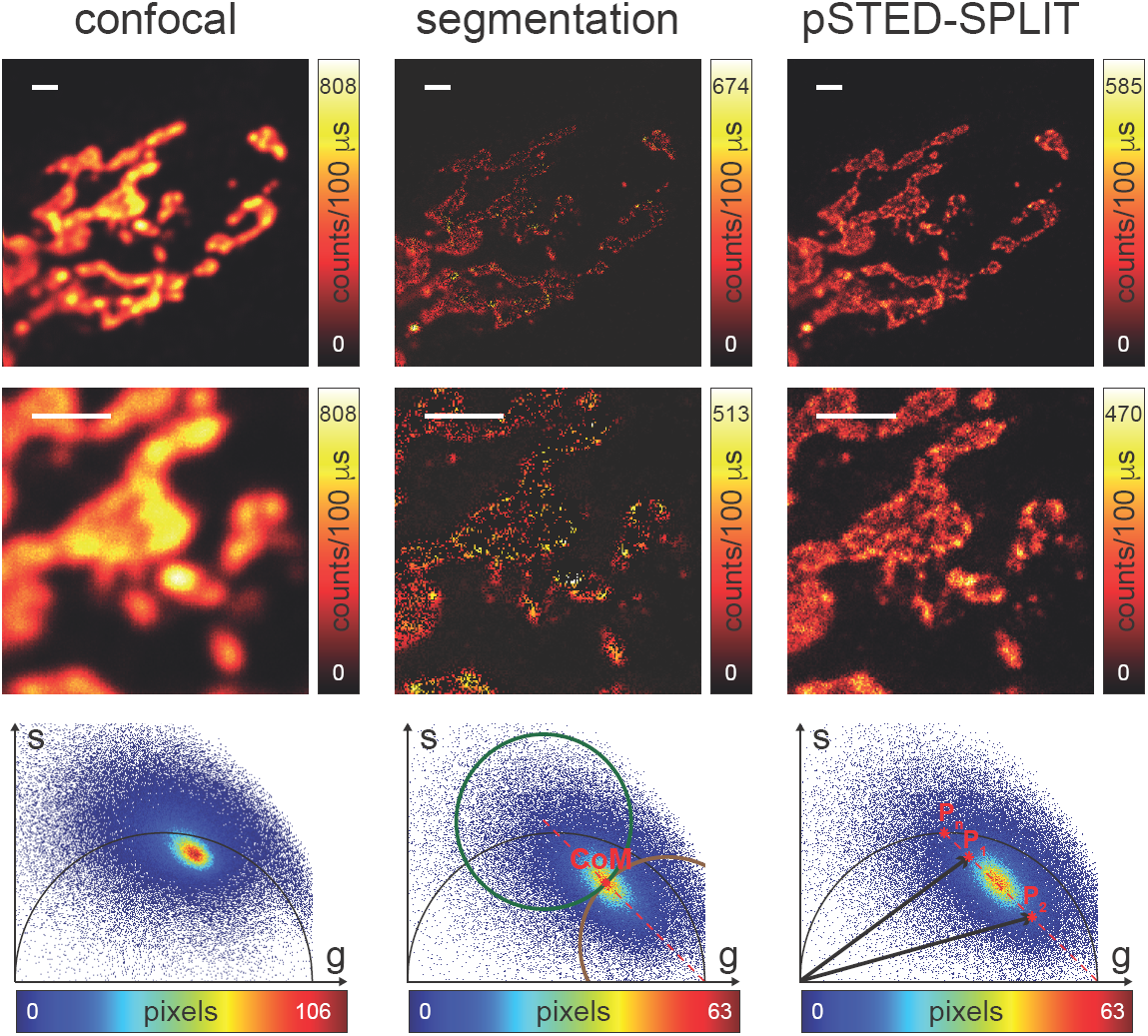
pSTED-SPLIT analysis of fixed cells. Confocal image (left) and the results of the pixel segmentation (center) and linear decomposition analysis (right) of time-resolved pSTED measurements of a sample of fixed cell with Atto 647N labeling mitochondria. Magnified views (central row) and phasor plot analysis (bottom) are also shown. Pixel-dwell time: 100 *μ*s. Pixel-size: 24 nm. Format: 512 *×* 512 pixels. Scale bars: 1 *μ*m.

## 4 Discussion

In this study we have extended the SPLIT technique introduced by us^16^ to the case of a pSTED implementation. We demonstrated with synthetic data and real data that the SPLIT-pSTED method is able to enhance the spatial resolution of a time-resolved pSTED measurement with no degradation on the SNR and with no artifacts introduction. The SPLIT-pSTED method rejects the photons emitted in the periphery of the excitation spot, *via* a simple linear decomposition algorithm in the phasors space, thus reducing the effective detection region without increasing the intensity of the STED beam and improving the spatial resolution. A comparison with a phasor-based segmentation methods is also carried out to demonstrate the higher robustness.

Thanks to the continuous emerging high photon collection efficiency (¿ 100 Mcps) time-resolved detection architecture,^17^ we envisage the integration of the SPLIT-STED method on any commercial STED microscope. Further studies will be concentrated in the application of the SPLIT approach in the context of point-scanning reversible saturable optical linear fluorescence transitions (RESOLFT) microscopy with reversible-switching fluorescent proteins (rsFP).^22^ Here, the rsFP’s fluorescence signal induced during RESOLFT illumination cycle (activation, de-activation and read-out), similar to STED microscopy, shows dynamics which change as distance of the rsdFPs from the center of the foci. Such spatial information encoded in the fluorescence dynamics can be decoded using the SPLIT approach to further enhance the resolution of the RESOFT images.

